# Mechanisms of sustaining oxygen extraction efficiency in dragonfly nymphs during aquatic hypoxia

**DOI:** 10.1101/2024.06.12.598728

**Authors:** Daniel J. Lee, Philip G.D. Matthews

## Abstract

Despite breathing water using their tidally ventilated rectal gills, dragonfly nymphs show a surprising ability to maintain oxygen (O_2_) extraction from the water during hypoxia. However, an increase in convective O_2_ transfer is insufficient to sustain aerobic demands by itself, which suggests that diffusive mechanisms must also be involved. This study examines the contributions of changing the O_2_ partial pressure gradient (PO_2_) and/or O_2_ conductance across the rectal gill in maintaining O_2_ extraction efficiency (OEE) of dragonfly nymphs during hypoxia. Data were collected using the same custom-designed respiro-spirometer described in a previous study with the addition of an implanted O_2_ sensor to measure hemolymph PO_2_. Results show that the implantation of the O_2_ sensor does not affect the respiratory and ventilatory response of nymphs to hypoxia. Hemolymph PO_2_ fell from 6.3 ± 1.6 kPa at normoxia to 2.5 ± 0.6 kPa at 16.0 kPa, which resulted in the PO_2_ diffusion gradient remaining statistically constant at these two water PO_2_s (17.5 ± 1.7 and 15.4 ± 0.7 kPa during normoxia and 16.0 kPa respectively). Beyond 16.0 kPa, a progressive reduction in hemolymph PO_2_ was unable to sustain the diffusion gradient. Mathematical modeling revealed that while the addition of hemolymph PO_2_ in tandem with ventilation frequency was able to elevate OEE during 16.0 kPa to that of normoxia, both were still insufficient during severe hypoxia and required an increase in O_2_ conductance. Estimating the change in whole-gill conductance showed that nymphs are indeed increasing their conductance as the water becomes hypoxic, demonstrating a reliance on both diffusion gradient and O_2_ conductance to enhance diffusive O_2_ transfer in conjunction with convective mechanisms to maintain O_2_ extraction during hypoxia.

## Introduction

Water is an energetically expensive medium to move due to its very high viscosity and density relative to air, and is also a poor source of oxygen (O_2_) due to its very low O_2_ capacitance coefficient (Brainerd and Ferry-Graham, 2005; Dejours, 1989; Maina, 2002; Ultsch, 1996). As such, all water-breathing animals face the innate challenge of having to expend a substantial portion of their total energy ventilating large volumes of an O_2_-deficient medium to satisfy their metabolic demands. To counter the above issues, the vast majority of water-breathers have evolved a unidirectional gas-exchange system, where water is continuously ventilated past a gas-exchange organ in a single direction. This constant unidirectional flow of water allows for energetically efficient ventilation as it requires less energy to keep this viscous and dense medium moving in a single direction (Brainerd and Ferry-Graham, 2005) while also allowing blood to establish a counter-current exchange system with the water by generating and maintaining a steep O_2_ partial pressure (PO_2_) diffusion gradient to enhance O_2_ uptake along the entire length of the exchange site (Rahn, 1966). Thus, unidirectional ventilation allows water-breathing animals to maximize O_2_ extraction while minimizing cost of breathing. From this perspective, aquatic dragonfly nymphs are an unusual exception as they breathe water using a tidally-ventilated rectal gill. Unlike unidirectional ventilation, tidal ventilation increases the cost of breathing as water flow must be repeatedly reversed in and out of the same opening (Brainerd and Ferry-Graham, 2005), and lowers O_2_ extraction since the inability of blood and water to form a counter-current system means that the PO_2_ gradient between these two phases diminishes over the course of a breath, reducing O_2_ diffusion (Piiper and Scheid, 1972). In addition, tidal ventilation is accompanied by the inevitable formation of an anatomical dead space, where a portion of the inhaled water is effectively trapped in the conducting water pathways and cannot exchange gases with the blood (Hughes and Morgan, 1973; Maina, 2002; Robertson, 2015). Thus, tidal ventilation greatly exacerbates the challenges of breathing water and one would predict that dragonfly nymphs would have reduced O_2_ extraction capabilities compared with unidirectional water-breathers. However, the study by Lee and Matthews (2024) demonstrated otherwise. Dragonfly nymphs exposed to hypoxic water were able to sustain both their overall metabolic rates and O_2_ extraction efficiency (ratio of O_2_ extracted to O_2_ inhaled; OEE) down to 5.3 kPa, demonstrating their ability to oxy-regulate even during severe O_2_ deprivation. When compared with other water-breathers, the dragonfly nymphs’ capability to maintain metabolic rate and OEE was not only on par with that of unidirectional-breathers, but also out-performed other tidal-breathers (Lee and Matthews, 2024), providing strong evidence that tidal ventilation in dragonfly nymphs is not associated with reduced O_2_ extraction during both normoxia and hypoxia. This is not the only unusual aspect of their gas-exchange strategy. Given the energetic challenges associated with transporting a medium as viscous and dense as water (Brainerd and Ferry-Graham, 2005; Maina, 2002), most water-breathing animals increase ventilatory flow primarily by increasing stroke/tidal volume, rather than ventilation frequency (Hughes and Saunders, 1970; Perry et al., 2009; Porteus et al., 2011; Rantin et al., 1992; Scott et al., 2008; Thomas and Gilmour, 2012). However, dragonfly nymphs oppose this trend as they increase gill minute ventilation exclusively by increasing ventilation frequency. Using iterative models to further investigate the effects of varying ventilation frequency and tidal volume on the nymphs’ OEE revealed that the same minute ventilation produced by manipulating either of these two parameters yielded the same calculated OEE. However, unlike the empirical data, these models showed progressive reductions in O_2_ extraction during hypoxia (Lee and Matthews, 2024). Thus, the observation that dragonfly nymphs seemingly maintain metabolic rate and OEE through just ventilation frequency is incomplete, and the authors proposed two additional unobserved factors that must also be changing in the animals.

The transfer of a gas between two compartments is driven by two processes: convection and diffusion. For ventilating animals, O_2_ is first convected from the surrounding environment into the gas-exchange organ, then passively diffuses through the gas-exchange surface and into the animal. Thus, both processes play critical roles and the transport of O_2_ by both convection and diffusion must exist in equilibrium to ensure steady transfer of O_2_. As the animal inhales the respiratory medium, dissolved O_2_ in water enters the gas-exchange organ as determined by:

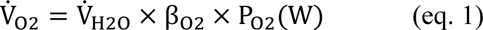

where V̇_O2_ is the O_2_ transfer rate, V̇_H2O_ is the ventilatory water flow (= minute ventilation), β_O2_ is the capacitance coefficient of O_2_ in water, and P_O2_(W) is the PO_2_ in water (Weibel, 1984). Of these three components of convective O_2_ transfer, β_O2_ is medium-specific while P_O2_(W) is determined by the quantity of O_2_ dissolved in water. Thus, they cannot be altered by the animal and any modulation of O_2_ transfer during convection must be accomplished through changes in the minute ventilation. Dragonfly nymphs show significant increases in their gill minute ventilation during hypoxia (Lee and Matthews, 2024), which indicates that V̇_H2O_ is indeed being elevated to counter the reduction in PO_2_(W) and maintain convective V̇_O2_. However, the observation that higher V̇_H2O_ by itself is unable to maintain overall metabolic rate and OEE means that diffusive V̇_O2_, which was not compensated for in the model, decreased as a consequence of the diminished PO_2_(W) and resulted in a progressive fall in overall O_2_ flux. Thus, nymphs must be employing mechanisms to maintain diffusive O_2_ transfer during hypoxia. The diffusion of O_2_ from the inhaled medium across the gas-exchange surface and into an animal is determined by:

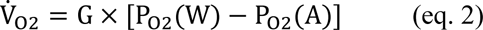

where G is the O_2_ conductance of the gas-exchange surface, and PO_2_(A) is the PO_2_ in the animal (Weibel, 1984). For dragonfly nymphs, G is that of the rectal gill and PO_2_(A) is the O_2_ tension in the tracheal system. G is further defined by:

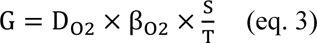

where D_O2_ is the diffusion coefficient of O_2_ through the gill epithelium, β_O2_ is the capacitance coefficient of O_2_ in tissue (assumed equal to β_O2_ of water), S is the surface area of the rectal gill, and T is the thickness of the diffusion barrier (Weibel, 1984). As such, the diffusive V̇_O2_ is determined by the O_2_ conductance of the gill and the PO_2_ diffusion gradient across it. For conductance, both D_O2_ and β_O2_ are material-specific values that cannot be modulated by the animal, and dragonfly nymphs are unlikely to be able to alter the surface area of their gill during acute hypoxia. The diffusion barrier thickness is largely comprised of the physical thickness of the rectal gill lamellae which is also unlikely to be changed by the animal during acute hypoxia. However, aquatic animals must contend with an additional component of the diffusion pathway: a stagnant boundary layer of water that adhering to the gas-exchange surface. This boundary layer further increases the thickness of the overall diffusion pathway, but unlike the rectal gill, its thickness can be reduced by increasing volumetric water flow moving past the gill surface (Pinder and Feder, 1990). Given that dragonfly nymphs substantially increase their gill minute ventilation during hypoxia, the boundary layer thickness must indeed be changing and providing an avenue for these insects to change O_2_ conductance across their gill. Dragonfly nymphs may also change the PO_2_ gradient by lowering the O_2_ tension in their tracheal system. This would potentially counter the reduction in water PO_2_ by allowing them to sustain a sufficiently large diffusion gradient to maintain O_2_ uptake from the hypoxic water. Thus, the compensation of reduced PO_2_ in the water to maintain passive O_2_ flux during hypoxia in dragonfly nymphs must be accompanied by changes in O_2_ conductance and/or the PO_2_ diffusion gradient.

The objective of this study was to investigate whether O_2_ conductance and/or PO_2_ diffusion gradient is changing in dragonfly nymphs during progressive hypoxia then determine the relative contributions of each parameter in sustaining OEE. The same respiro-spirometer from the previous study (Lee and Matthews, 2024) was used to obtain the ventilatory and respiratory parameters, except this time, the nymphs were also implanted with an O_2_ sensor to directly measure hemolymph PO_2_ (proxy for tracheal PO_2_). The resulting tidal volume, ventilation frequency, metabolic rate, and OEE were compared with those from the previous study to assess the effect of implantation on the dragonfly nymphs’ response to hypoxia, then the model simulation was used to calculate the simulated OEE under three scenarios: 1) ventilation frequency, 2) ventilation frequency and PO_2_ diffusion gradient, and 3) ventilation frequency, PO_2_ diffusion gradient, and O_2_ conductance. Dragonfly nymphs are predicted to change both their O_2_ conductance and PO_2_ diffusion gradient in order to maintain O_2_ uptake during hypoxia, as the innate difficulty with extracting O_2_ from water likely demands both strategies to compensate for reduced O_2_ availability.

## Materials and methods

### Animals

Early-final instar *Anax junius* (Drury, 1773) nymphs were captured, identified, and treated as described previously between May and August 2024 (Lee and Matthews, 2024). Briefly, nymphs were captured from the South campus research ponds at the University of British Columbia Point Grey campus and individually housed in 1 l glass aquaria connected to a central re-circulating system. Dechlorinated Vancouver tap water was used in all experiments.

### Measurement of ventilatory and respiratory parameters

In order to measure the tidal volume, breathing frequency, metabolic rate, and hemolymph PO_2_ of nymphs during various water PO_2_ levels, the same setup and strategy was used as described previously (Lee and Matthews, 2024), except for the following changes: Once the respiro-spirometer was fully assembled with the nymph and all connections, the syringe barrel reservoir was bubbled with 21.3 kPa PO_2_ balance N_2_ gas mixture and the peristaltic pump was started to continuously re-circulate the water through the respiration/ventilation chamber. Then the water in the outer experimental chamber was removed until the water level was below the ventral surface of the nymph. This exposed the portion of the animal anterior to the rear harness out of the water while still allowing the nymph to ventilate oxygenated water through its rectum. The dorsal thoracic surface of the nymph was dried with paper towels, and a calibrated O_2_ sensor was implanted into the hemocoel of the animal as described previously (Lee et al., 2018). Once the vinyl polysiloxane (VPS) sealing the implanted sensor tip and the hole in the nymph’s tergite cured, the experimental chamber was re-filled with water and the nymph was allowed to acclimate to the setup for at least 12 h overnight (21.3 kPa PO_2_ and 4 ml/min water flow rate). The data collection followed an identical procedure as in the previous study (Lee and Matthews, 2024), with the only change being the addition of the implanted O_2_ sensor which was recorded simultaneously with the other data.

### Data analysis

Running the chamber without a nymph as a ‘blank’ to check for offset between the in-current and ex-current O_2_ sensors showed an offset ranging from 3.3 to 6.3% air-sat (0.7 to 1.3 kPa) in normoxia and −1.8 to 1.4% air-sat (−0.4 to 0.3 kPa) in 5.3 kPa. Although the same trend of the offset decreasing towards lower PO_2_ was observed here, the magnitude of the range was noticeably higher than that seen previously (Lee and Matthews, 2024). Thus, it was determined that sensor offset may have noticeable effects during data analysis even at lower PO_2_ levels and so the offset was corrected at all exposure PO_2_s as follows: For each nymph, the O_2_ data recorded at each exposure PO_2_ (21.3, 16.0, 10.7, and 5.3 kPa) was screened for a breath-hold period, which revealed that four out of seven nymphs had breath-hold periods during at least two exposure PO_2_s. For these four individuals, the average ex-current PO_2_ during breath-hold was subtracted from the corresponding average in-current PO_2_ to directly calculate the sensor offset at that exposure PO_2_. Then, a linear regression relating in-current PO_2_ to calculated sensor offset was plotted, and was subsequently used to estimate sensor offset during the other experimental PO_2_s with no breath-hold period. Of the remaining three nymphs, two showed breath-hold periods only at one exposure PO_2_, and as a result, the blank run offset at 5.3 kPa was used together with the exposure PO_2_ offset to create the linear curve. One nymph did not show any breath-hold periods during the entire experiment and could not be corrected. However, comparing the data of this nymph to those of corrected nymphs showed that the uncorrected nymph lies well within range of the others and did not present an outlier. Thus, it was included in the final data analysis and the data left uncorrected.

Analysis of tidal volume, breathing frequency, gill minute ventilation, WCR, and OEE followed the same procedure as described previously (Lee and Matthews, 2024). For metabolic rate, the only change was including the Δ_Offset_ parameter for all exposure PO_2_s rather than at just normoxia. In order to calculate hemolymph PO_2_ and diffusion gradient, the hemolymph PO_2_ data was screened to find the same time section as the ventilation data then averaged to find the mean hemolymph PO_2_. This mean hemolymph value was subtracted from the mean in-current PO_2_ to finally calculate the diffusion gradient between the insect and the surrounding water, and was repeated for all exposure PO_2_s.

Data were analyzed in R v4.2.2 (R Core Team, 2022) and all data were analyzed using one-way repeated measures ANOVA (nlme package v3.1-160) to test for any statistical differences. Data are shown as means ± s.e.m unless otherwise stated.

## Results

### Controlled PO_2_ versus measured in-current PO_2_

As was observed previously (Lee and Matthews, 2024), the measured in-current PO_2_ was slightly elevated relative to the controlled levels in this study (Table 1). This is most likely due to the same reason of O_2_ super-saturation during normoxia and passive influx from the surrounding air during hypoxia, and thus the same data and statistical analysis strategy was used here as described previously (Lee and Matthews, 2024): The measured PO_2_ values were used for data analysis while the controlled PO_2_ were used for statistical analysis and visual presentation.

**Table 1.** Comparison of mass-flow controlled PO_2_ to measured PO_2_ at each O_2_ level.

| **% air saturation** | **Controlled PO<sub>2</sub> (kPa)** | **Measured PO<sub>2</sub> (kPa)** | **% difference** |
| --- | --- | --- | --- |
| 100 | 21.3 | 23.8 | 12 |
| 75 | 16.0 | 17.9 | 12 |
| 50 | 10.7 | 12.0 | 12 |
| 25 | 5.3 | 6.2 | 18 |

### Ventilatory and respiratory parameters during implantation

The mean breathing frequency, tidal volume, gill minute ventilation, and metabolic rate were measured from seven implanted early-final *A. junius* nymphs (mean weight 1.2 ± 0.1 g). At 21.3 kPa (normoxia), nymphs had a mean breathing frequency of 13.2 ± 3.1 BPM which was not significantly different from those at 16.0 and 10.7 kPa PO_2_ (13.9 ± 2.6 and 19.9 ± 2.7 BPM, respectively) (Fig. 1a). However, there was a significant increase in breathing frequency to 26.6 ± 1.9 BPM at 5.3 kPa, which was statistically the highest value (One-way repeated measures ANOVA, *F*=11.907, d.f=3, *P*<0.0002). The mean tidal volume was 57.6 ± 8.3 μl at normoxia, and although it increased to a maximum of 69.0 ± 11.8 μl at 10.7 kPa, tidal volume did not change significantly with PO_2_ (One-way repeated measures ANOVA, *F*=1.069, d.f=3, *P*=0.3868) (Fig 1b). The gill minute ventilation, a product of breathing frequency and tidal volume, thus changed in a similar manner as breathing frequency, with a mean value of 762.9 ± 267.5 μl H_2_O/min at normoxia that significantly increased to 1833.0 ± 392.6 μl H_2_O/min at 5.3 kPa (One-way repeated measures ANOVA, *F*=6.962, d.f=3, *P*<0.005) (Fig. 2a). Nymphs had a mean metabolic rate of 1.3 ± 0.4 μl O_2_/min at normoxia, and this did not significantly change during any of the hypoxia exposures (One-way repeated measures ANOVA, *F*=0.875, d.f=3, *P*=0.472) (Fig. 2b). The WCR of nymphs at normoxia was 612.9 ± 97.4 μl H_2_O/μl O_2_ which was not significantly different from the values at 16.0 and 10.7 kPa (934.5 ± 111.3 and 1564.4 ± 337.1 μl H_2_O/μl O_2_). At 5.3 kPa, however, there was a significant increase to 2546.9 ± 790.5 μl H_2_O/μl O_2_ (One-way repeated measures ANOVA, *F*=4.877, d.f=3, *P*<0.02) (Fig 3a). Nymphs had a resting OEE of 30.6 ± 5.7% at normoxia, and maintained this value during all hypoxia exposures (One-way repeated measures ANOVA, *F*=1.021, d.f=3, *P*=0.407) (Fig. 3b).

**Fig. 1.**
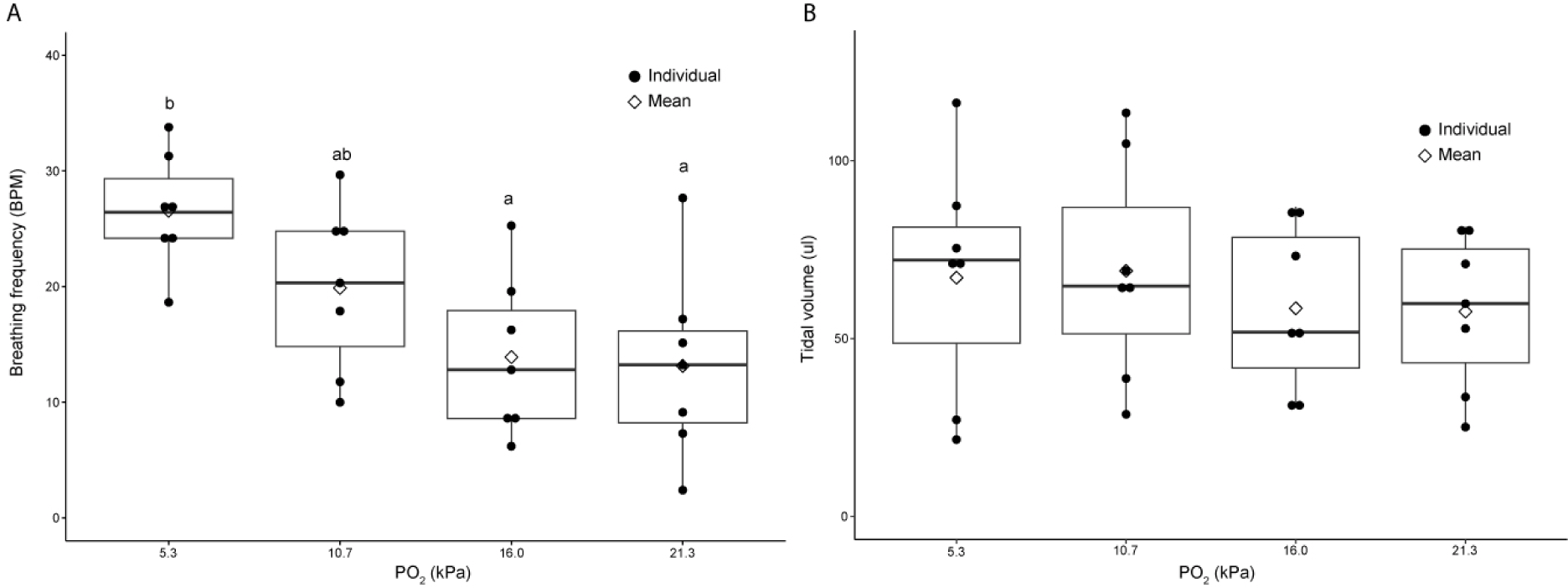
Changes in ventilation frequency (A) and tidal volume (B) of early-final instar *Anax junius* nymphs (n=7) across different PO2 levels, shown as boxplots. Filled symbols represent individual data, while blank symbols represent mean values. Letters represent significant differences where applicable.

**Fig. 2.**
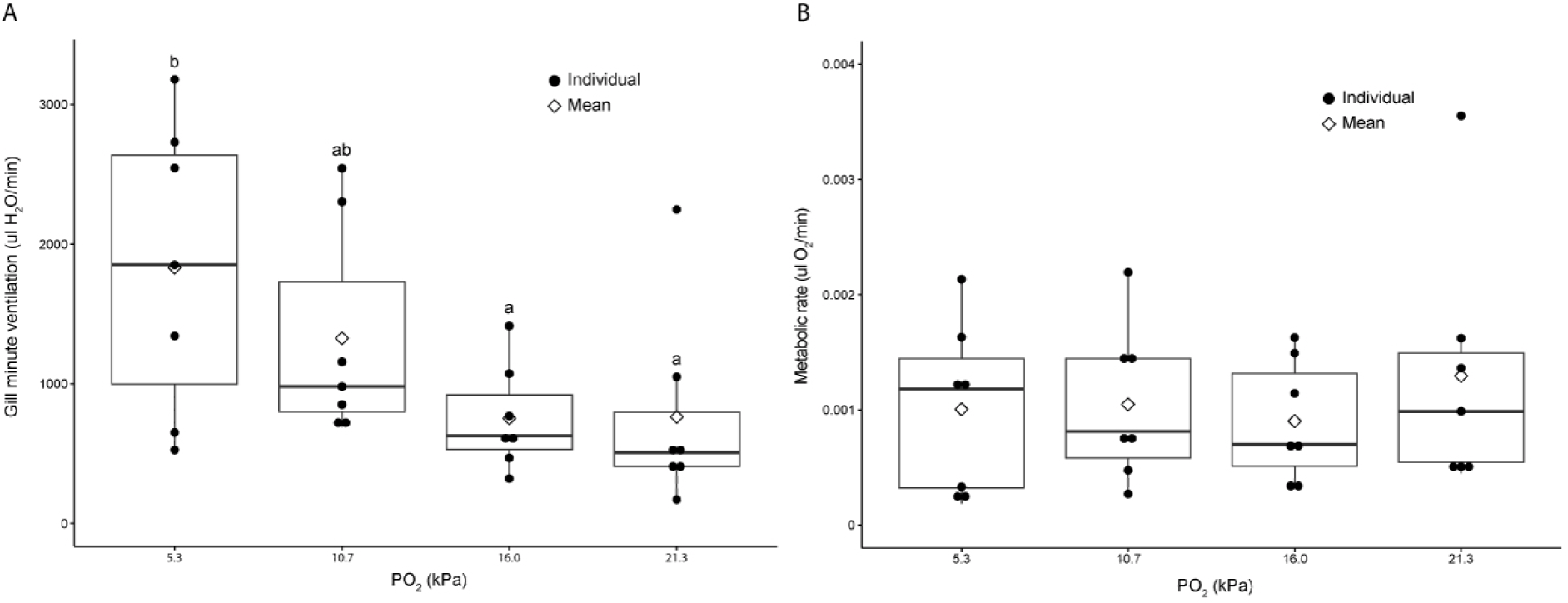
Changes in gill minute ventilation (A) and metabolic rate (B) of early-final instar *Anax junius* nymphs (n=7) across different PO2 levels, shown as boxplots. Filled symbols represent individual data, while blank symbols represent mean values. Letters represent significant differences where applicable.

**Fig. 3.**
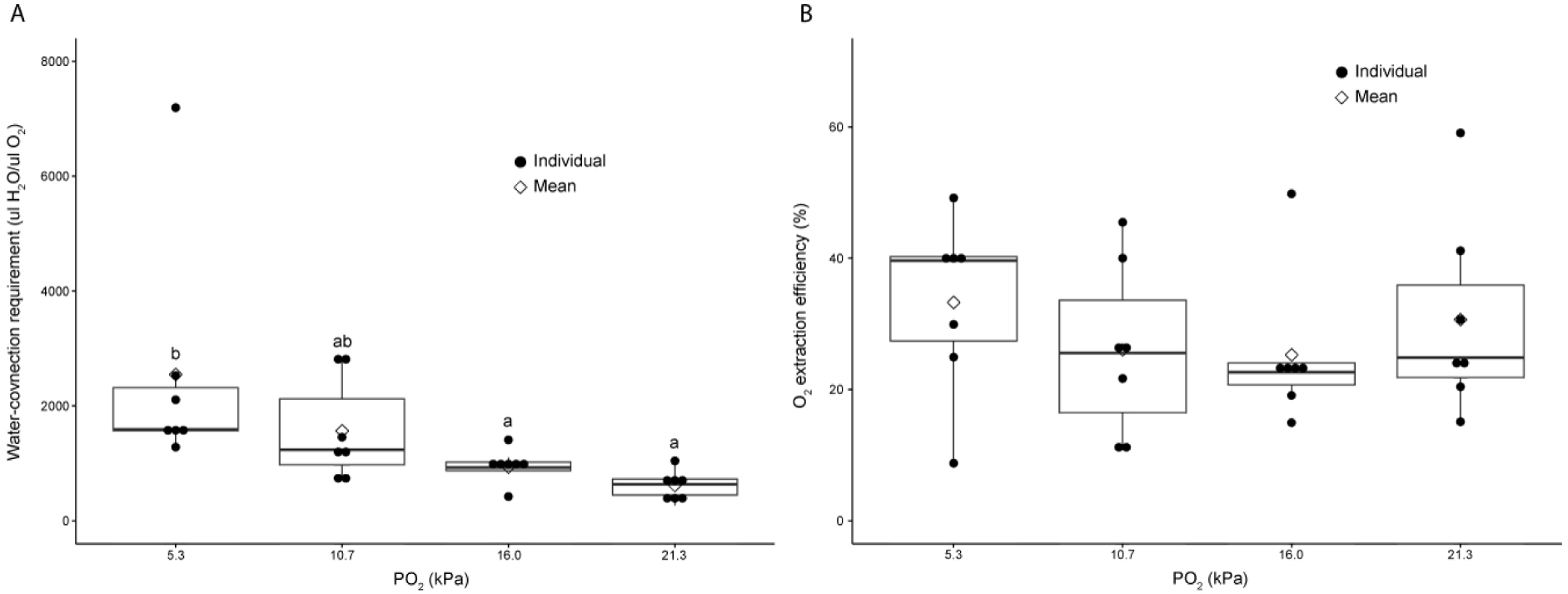
Changes in water-convection requirement (A) and oxygen extraction efficiency (B) of early-final instar *Anax junius* nymphs (n=7) across different PO2 levels, shown as boxplots. Filled symbols represent individual data, while blank symbols represent mean values. Letters represent significant differences where applicable.

### *In vivo* hemolymph PO_2_ and diffusion gradient

At normoxia, *A. junius* nymphs had the highest hemolymph PO_2_ of 6.3 ± 1.6 kPa, which was significantly reduced to 2.5 ± 0.6 kPa at 16.0 kPa PO_2_ (Fig. 4). As the water PO_2_ decreased further, there were consistent reductions in hemolymph PO_2_ (2.0 ± 0.8 and 0.5 ± 0.2 kPa at 10.7 and 5.3 kPa, respectively), however these changes were not significantly different from the value at 16.0 kPa (One-way repeated measures ANOVA, *F*=11.471, d.f=3, *P*<0.0005). The PO_2_ diffusion gradient between the nymphs and surrounding water was 17.5 ± 1.7 kPa at normoxia, and nymphs maintained this value at 16.0 kPa (15.4 ± 0.7 kPa) (Fig. 4). However, the diffusion gradient was significantly reduced at lower water PO_2_ levels, with 5.3 kPa having statistically the lowest diffusion gradient of 5.7 ± 0.1 kPa (One-way repeated measures ANOVA, *F*= 53.293, d.f=3, *P*<0.0001).

**Fig. 4.**
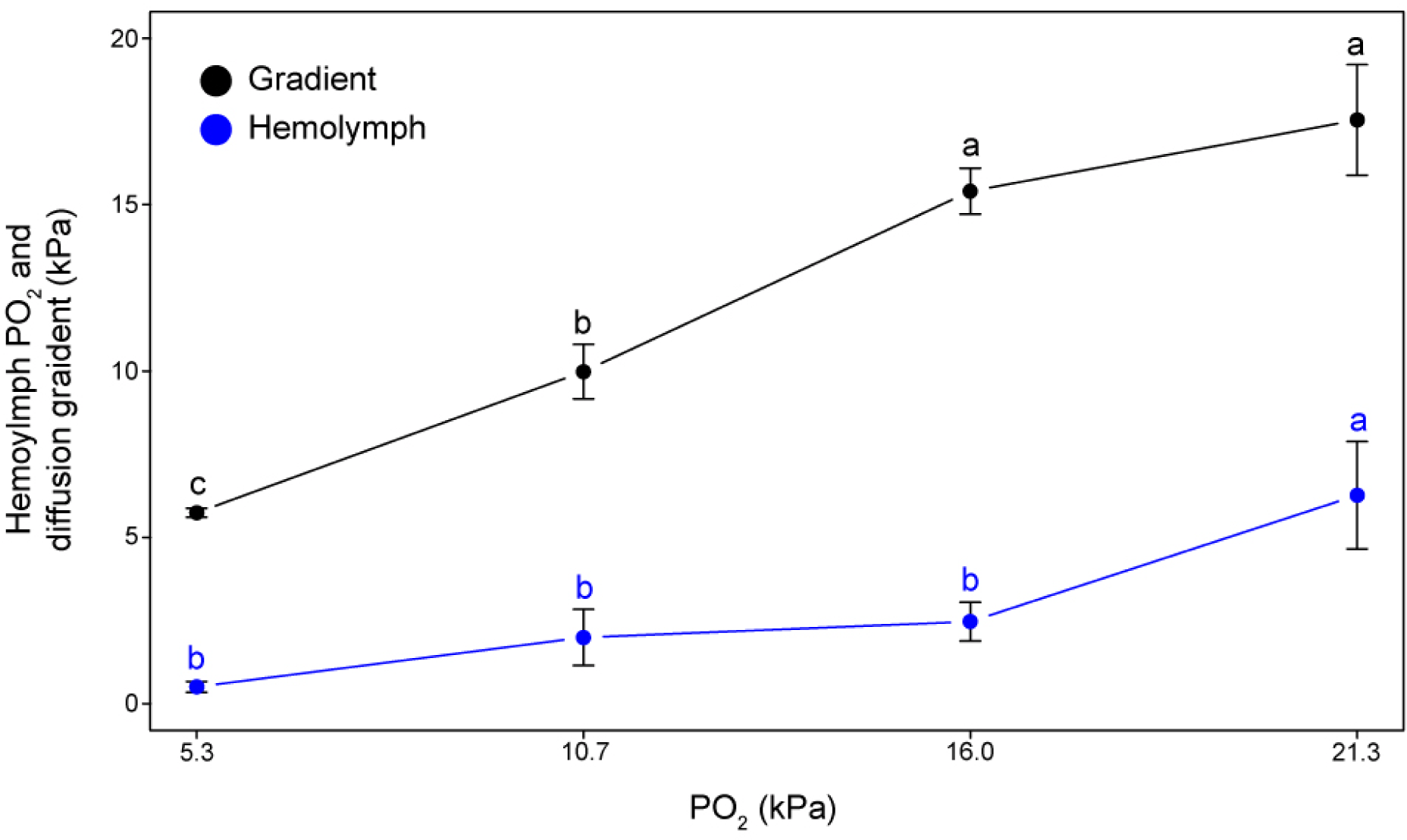
Changes in hemolymph PO_2_ (blue) and PO_2_ diffusion gradient between the insect and the water (black) of early-final *Anax junius* nymphs (n=7) across different PO_2_ levels. Letters represent significant differences where applicable.

## Discussion

### Ventilatory and respiratory parameters of implanted nymphs

Comparing the ventilatory and respiratory data of implanted nymphs to those from the previous study (Lee and Matthews, 2024) shows that the two data sets are in excellent agreement. The resting tidal volume at normoxia differed only by 3 μl (57.6 versus 60.8 μl) and there were no significant changes during hypoxia in either group of animals. As a result, the hypoxic ventilatory response in both groups exclusively involves an increase ventilation frequency. Ventilation frequency appears slightly elevated at normoxia during implantation, as it was statistically the same as those during 16.0 and 10.7 kPa which indicates that normoxic nymphs are breathing as fast as those that are exposed to moderate hypoxia. While this may be a different trend from non-implanted nymphs which show a significant increase in ventilation frequency from normoxia to 10.7 kPa (Lee and Matthews, 2024), the same overall numerical and statistical patterns are observed in both studies: breathing frequency increases with hypoxia until reaching statistically the highest value at 5.3 kPa. The elevated breathing frequency of implanted nymphs at normoxia is likely due to sampling variation rather than a direct stress response to implantation, as comparing the metabolic rates of implanted and non-implanted nymphs shows 1) nearly identical values at normoxia (1.3 versus 1.4 μl O_2_ min^-1^) and 2) no significant change in metabolic rate even during severe hypoxia in either group. That the implantation did not elicit visible differences in metabolic rate provides strong evidence that nymphs were able to assume ventilatory and respiratory patterns that are indistinguishable from the non-implanted animals in the previous study (Lee and Matthews, 2024). Perhaps the most noticeable difference between implanted and non-implanted animals is the OEE, where implanted nymphs had a 25% lower value during normoxia (30.6 versus 40.1%). However, given that the underlying ventilatory (tidal volume, breathing frequency) and respiratory (metabolic rate) parameters are very similar and both groups show the same statistical trend, it is again likely a product of sampling variation rather than an experimentally-induced difference.

Due to their closed tracheal system which prevents entry into the gas-conducting vessels, it was not possible to measure the dragonfly’s tracheal PO_2_ in this experiment. Instead, their hemolymph PO_2_ tensions were measured as a proxy, with the assumption that the two compartments will likely be in equilibrium due to the open circulatory system and constant stirring of hemolymph due to abdominal pumping during ventilation. At first glance, the observation that the hemolymph PO_2_ of dragonfly nymphs during normoxia is just 30% of water saturation levels is not surprising given the respiratory challenges of breathing water. The combination of innately high ventilation cost and low O_2_ concentration in water (Brainerd and Ferry-Graham, 2005; Dejours, 1989; Maina, 2002; Ultsch, 1996) with the consequences of tidal ventilation (Brainerd and Ferry-Graham, 2005; Piiper and Scheid, 1972) necessitates that dragonfly nymphs maximize O_2_ extraction from the water, which is likely achieved in part by maintaining a steep PO_2_ diffusion gradient from water to a low hemolymph PO_2_. However, comparing the dragonfly nymphs in the current study with the only other known study of internal PO_2_ in aquatic insects (Birrell et al., 2023 preprint) reveals a stark contrast. While the mean internal PO_2_ of dragonfly nymphs matched remarkably well in both studies (6.3 kPa versus 5.9 kPa), they are noticeably higher than those of other water-breathing insects (mean 0.9 kPa; Birrell et al., 2023 preprint). Why dragonfly nymphs have a relatively elevated internal PO_2_ is unknown. However, a potential explanation may be associated with the ability of dragonfly nymphs to alter their hemolymph PO_2_ as a mechanism to sustain O_2_ uptake during hypoxia. While an elevated internal PO_2_ may reduce O_2_ extraction during normoxia due to a lesser PO_2_ diffusion gradient, it also provides an avenue for the insect to compensate for reduced O_2_ availability without having to alter gill conductance and gill minute ventilation. This is precisely what is observed from the dragonflies: a significant reduction in hemolymph PO_2_ at 16 kPa water PO_2_ maintains the diffusion gradient between the gill water and the nymph’s hemolymph (Fig. 4) which enables them to maintain their metabolic rate without a change in gill minute ventilation (Figs. 2). A survey of the literature reveals that unlike dragonflies, most water-breathing insects (including members of lineages studied by Birrell et al.) are unable to maintain their metabolic rates as the water becomes hypoxic (e.g. Bäumer et al., 2000; Brodersen et al., 2004; Galic et al., 2019; Malison et al., 2020a; Malison et al., 2020b). This progressive decline in metabolic rate, in spite of a potential change in gill conductance and/or gill ventilation, may indeed be partly due to these insects maintaining an extremely low internal PO_2_ even during normoxia. As a result, they would have no capacity to sustain or increase their water-to-hemolymph PO_2_ diffusion gradient during hypoxia.

### Contributions of ventilation frequency, diffusion gradient, and O_2_ conductance to OEE

To estimate the contributions of ventilation frequency, PO_2_ diffusion gradient, and O_2_ conductance to OEE, the same mathematical model introduced in Lee and Matthews (2024) was used in this study and is described in greater detail to highlight the mathematical relationships and equations used to simulate O_2_ transfer between the surrounding water and the insect during ventilation. The model simulates the incremental O_2_ extraction during inhalation and O_2_ loss during exhalation in the dragonfly nymph through the application of Fick’s principle of gas transfer:

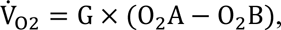

whe re V̇_O2_ is O_2_ transfer rate, G is conductance, and O_2_A and O_2_B is the O_2_ content in compartments A and B.

The model first considers the bulk flow of O_2_ within the water inhaled into the branchial chamber as determined by:

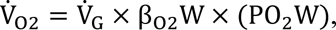

whe re V̇_G_ is gill minute ventilation, β_O2_W is capacitance coefficient of O_2_ in water, and PO_2_W is the PO_2_ in the inhaled water. For each time increment, the amount of water inhaled into the branchial chamber is calculated as:

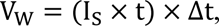

where V_W_ is the volume of water inhaled (μl), I_S_ is the inhalation rate slope (μl s^-2^), t is the time since start of inhalation (s), and Δt is the time increment (s). The product of I_S_ and t yields the instantaneous inhalation rate (μl s^-1^) at that specific time since start of inhalation, and the product of the instantaneous inhalation rate and Δt determines how long (and therefore how much) the nymph inhaled. Then, V_W_ is multiplied by β_O2_ and PO_2_W to determine the amount of O_2_ (μl) that has been inhaled into the branchial chamber at that specific time point. This newly introduced water and O_2_ volume is added to any pre-existing water and O_2_ in the branchial chamber, and the final PO_2_ in the branchial chamber is found as:

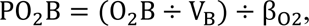

where O_2_B is branchial O_2_ (μl) and V_B_ is branchial water (μl). At this point, the gas-exchange switches to passive diffusion from the branchial chamber into the tracheal system of the insect, using the equation:

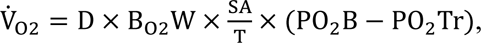

where D is diffusion coefficient of O_2_ in water, SA/T is surface area to thickness ratio of the rectal gill, and PO_2_Tr is PO_2_ in the tracheal system. However, SA/T of dragonfly nymph rectal gills is unknown, and as such D, B_O2_, and SA/T were collectively treated as the single O_2_ conductance value G (μl O_2_ min^-1^ kPa^-1^) which was determined as described in the previous study (Lee and Matthews, 2024). The volume of new O_2_ diffusing across the rectal gill and into the tracheal system is calculated by:

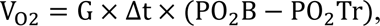

where Δt is time increment in minutes, then is added to the pre-existing volume of O_2_ in the tracheal system to find the current total volume of O_2_:

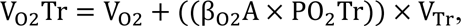

where β_O2_A is the capacitance coefficient of air (μl O_2_ μl H_2_O^-1^ kPa^-1^) and V_Tr_ is the volume of the tracheal system (μl). The tracheal system volume was estimated from micro-CT cross-sections as described previously (Lee and Matthews, 2024). The incremental consumption of O_2_ due to the metabolic rate of the animal is subtracted from the total tracheal O_2_ volume, and the remaining tracheal O_2_ volume is converted to a new PO_2_Tr to be used in the next inhalation increment. Lastly, the model calculates the volume of O_2_ left in the branchial chamber after diffusion into the animal to determine the pre-existing volume of O_2_ in the branchial chamber for the next inhalation increment. This iterative process is repeated until the nymph has inhaled its full tidal volume.

During exhalation, the above process is reversed. First, the metabolically consumed O_2_ volume is subtracted from the current O_2_ volume in the tracheal system, and the remaining V_O2_Tr is converted to PO_2_Tr. Then, a new V_O2_ diffuses from the branchial chamber into the tracheal system as determined by the diffusion gradient and G value, and the remaining O_2_B is divided by the current V_B_ to find the O_2_ concentration per μl of water. Using the exhalation rate slope, time since start of exhalation, and Δt, the volume of water exhaled by the animal is calculated for that specific time increment, and the corresponding volume of O_2_ exhaled is determined. Finally, the water and O_2_ volume remaining in the branchial chamber after the exhalation increment is used to calculate the remaining PO_2_ in the branchial chamber which determines the diffusion gradient for the next time increment. This iterative process is repeated until the nymph has exhaled its full tidal volume, then the cycle begins with the next inhalation. The end of each inhalation provides the starting conditions for each exhalation and vice versa. Simulated OEE was calculated by dividing the total O_2_ extracted into the tracheal system during both inhalation and exhalation by the total O_2_ inhaled.

Using the empirically measured ventilatory and respiratory parameters from the current study, the above mathematical model was used to estimate the contribution of ventilation frequency to OEE in the absence of changing hemolymph PO_2_ or O_2_ conductance by replicating the gill minute ventilation at 21.3, 16.0, 10.7, and 5.3 kPa PO_2_ via ventilation frequency (Model 1). In addition, the corresponding tidal volume and metabolic rate values measured from nymphs at each experimental PO_2_ were used in the models in order to simulate the nymphs’ ventilatory response as accurately as possible. Then, the *in vivo* hemolymph PO_2_ measured at each experimental PO_2_ was introduced into Model 1 to observe the impact of changing PO_2_ diffusion gradient on altering OEE (Model 2). Finally, the G value at 16.0, 10.7, and 5.3 kPa was manually adjusted until the simulated OEE in Model 2 matched the normoxic value (Model 3) to determine if the O_2_ conductance must change in addition to ventilation frequency and PO_2_ diffusion gradient. All measured and estimated parameters used for the model simulations are shown in Table 2.

**Table 2.**
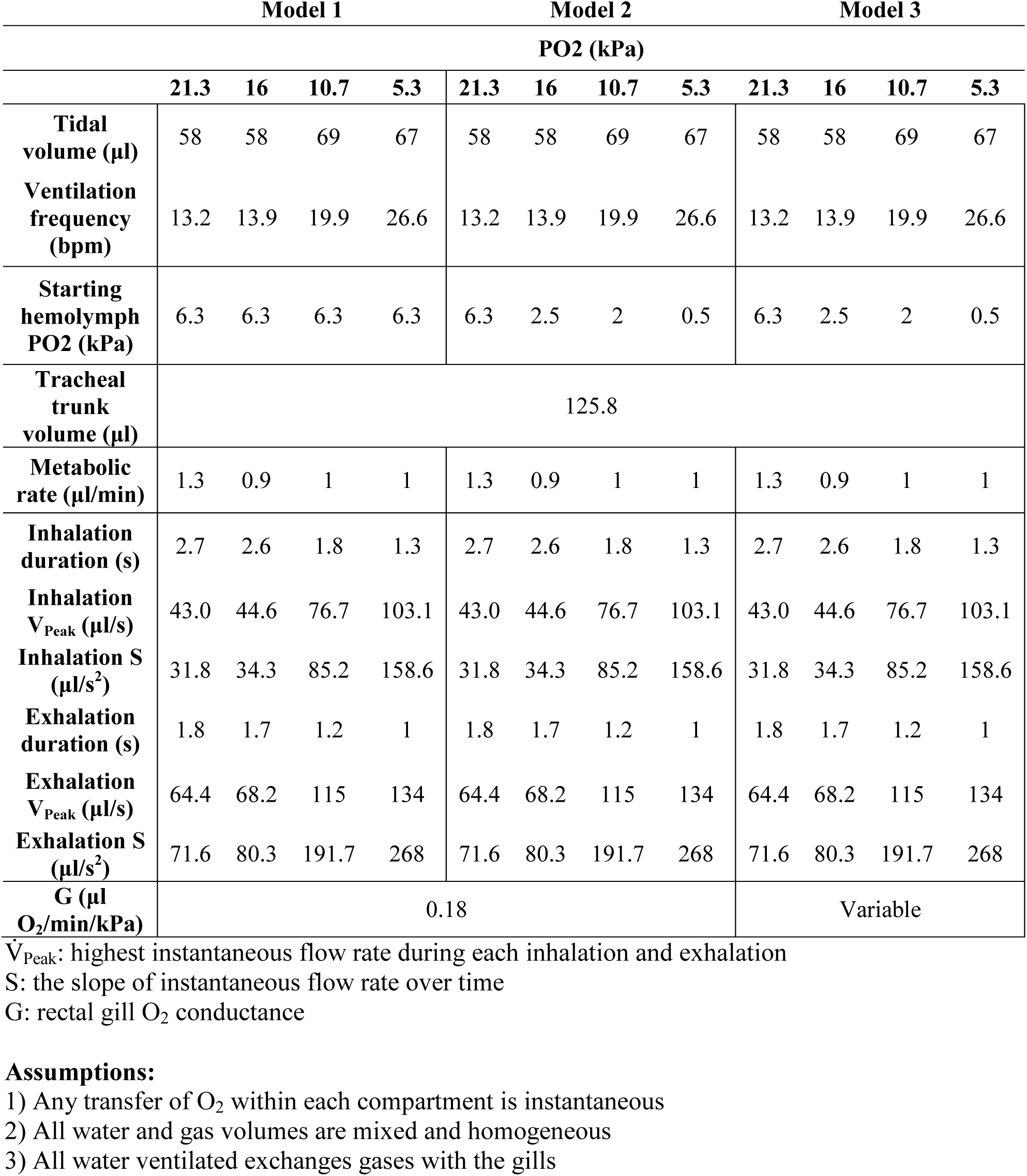
All parameters, conditions, and assumptions used for the theoretical modeling of dragonfly nymph oxygen extraction efficiency under three scenarios.

The observation that Model 1 simulated a progressive decrease in OEE with declining water PO_2_ (31, 25, 13, and 0% during 21.3, 16.0, 10.7, and 5.3 kPa, respectively) matches the simulation results from the previous study (Lee and Matthews, 2024), and confirms that ventilation frequency alone is insufficient to sustain OEE (Fig 5). Interestingly, the model resulted in a progressive loss of O_2_ from the animal into the water at 5.3 kPa, due to the fact that the starting hemolymph PO_2_ was higher than the surrounding water. Thus, it was determined that under the specified conditions of Model 1, the animal must change hemolymph PO_2_ or gill conductance in addition to ventilation frequency to extract O_2_ at 5.3 kPa, and the contribution of just ventilation frequency was set to zero. Incorporating hemolymph PO_2_ simultaneously with ventilation frequency (Model 2) caused a substantial increase in simulated OEE at all experimental PO_2_s, the most notable of which was at 16.0 kPa where OEE increased from 25 to 35%. This simulated ability of lowered hemolymph PO_2_ to elevate OEE back to normoxic levels is not surprising given the ventilatory and respiratory data observed in this study. Between 21.3 and 16.0 kPa, there were no significant differences in gill minute ventilation, PO_2_ diffusion gradient, or metabolic rate. This means that the overall conditions for O_2_ uptake in these two PO_2_ levels were the same, and by incorporating hemolymph PO_2_ into the calculation, Model 2 was able to recreate this scenario. However, the addition of variable hemolymph PO_2_ is still unable to elevate OEE to match normoxic values at the more severe hypoxic conditions of 10.7 and 5.3 kPa. Again, this is in excellent agreement with the empirically measured data. The inability of the animals to maintain their PO_2_ diffusion gradient during the above experimental conditions (Fig. 4) indicates a fall in overall O_2_ diffusion rates from the water into the animal and, if left uncompensated for as in Model 2, would predictably result in a reduction in OEE. Thus, the gill O_2_ conductance (G) between the insect and the water must also change in the model simulation in order to maintain OEE during severe hypoxia (Model 3; Fig. 5). The observation that G must change in addition to ventilation frequency and hemolymph PO_2_ in the model simulation raises the question: are dragonfly nymphs physiologically capable of altering the G of their rectal gill to enhance O_2_ extraction?

**Fig. 5.**
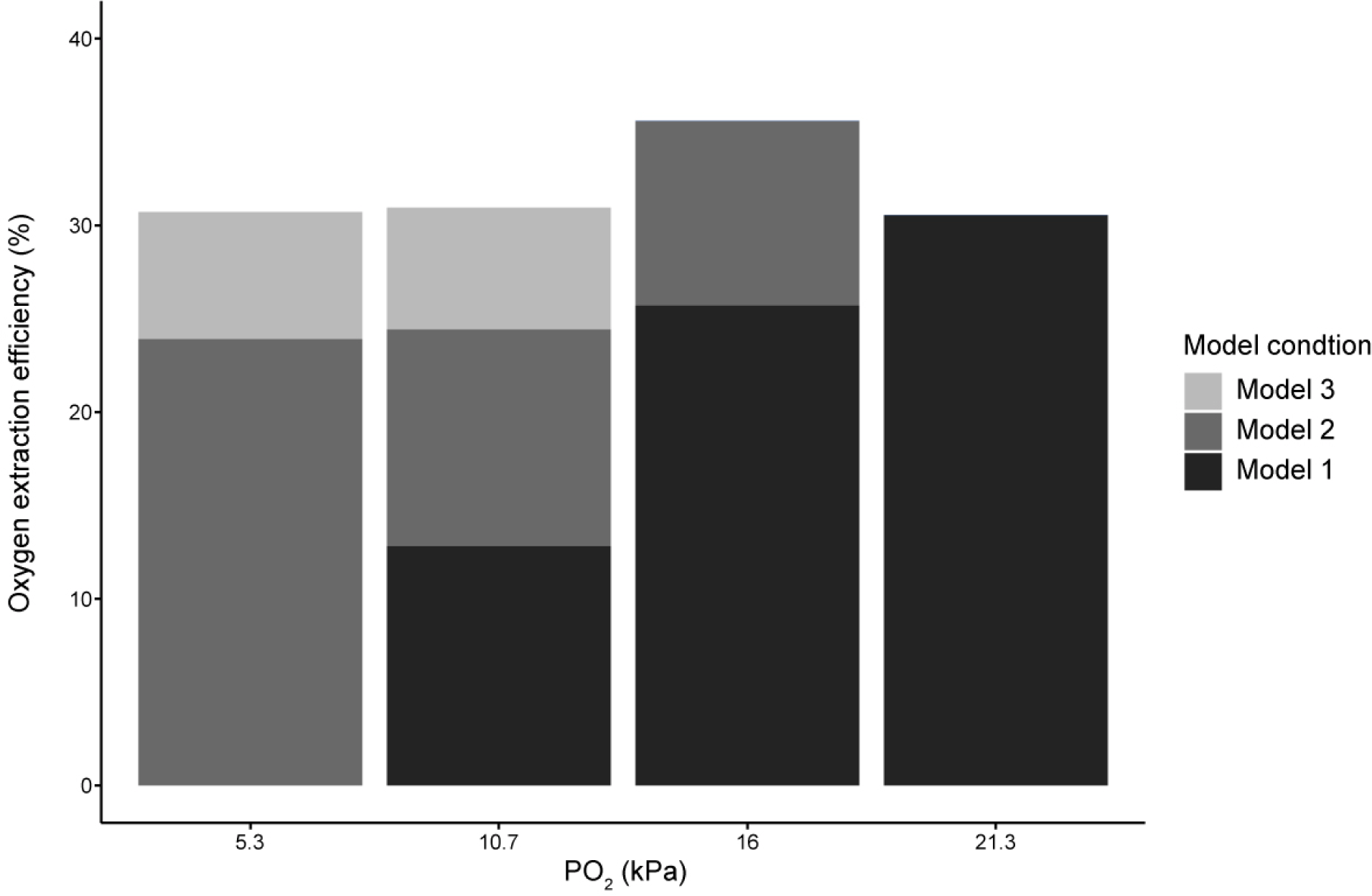
Summary of model simulated oxygen extraction efficiency results under three different conditions. Model 1: ventilation frequency only; Model 2: ventilation frequency and hemolymph PO_2_; Model 3: ventilation frequency, hemolymph PO_2_, and oxygen conductance.

### Modulation of gill conductance during hypoxia

Since the surface-area-to-thickness ratio is the only component of G that can be changed by the animal, an increase in G must correspond to an increase in the ratio. Assuming that the surface area of the rectal gill remains constant, this change in the area-to-thickness ratio must in turn be accomplished by a reduction in the overall diffusion barrier thickness. The physical gill epithelial thickness in dragonfly nymphs is unlikely to change during hypoxia, and so the only remaining avenue for increasing conductance would be to decrease the thickness of the stagnant boundary layer of water adhering to the gill’s surface. The mathematical relationship between boundary layer thickness and laminar water flow (Feder and Burggren, 1985) states that:

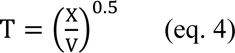

where T is boundary layer thickness, X is length of the gas exchange site, and V is water flow adjacent to the boundary layer. Assuming that X remains constant, the observed 2.4 fold increase in nymph gill minute ventilation during 5.3 kPa relative to normoxia would theoretically achieve a 35% reduction in T by itself. In addition, the increasingly turbulent tidal water flow during hyperventilation should further decrease boundary layer thickness beyond what is predicted based on laminar flow (Hitchman, 1978), thus presenting the increased ventilation frequency of the dragonfly nymphs as a substantial avenue for altering boundary layer thickness. Considering that the boundary layer can comprise between 80 to 90% of total diffusion resistance (Hills and Hughes, 1970), a reduction in boundary layer thickness should in turn greatly increase G between the insect and the water. Although the values of O_2_ conductance could not be empirically measured in the dragonfly nymphs, it is possible to estimate their whole-gill conductance for O_2_ during normoxia and hypoxia to test whether these insects do indeed increase their gill conductance during hypoxia. Rearranging Eq. 2 shows that G is defined as:

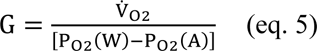

whe re V̇_O2_ is the O_2_ flux or metabolic rate, P_O2_(W) is the PO_2_ in the water, and P_O2_(A) is the PO_2_ in the hemolymph. Since all three parameters have been empirically measured during the experiment, it is then possible to calculate and see whether the physiological G changes from normoxia to hypoxia. Computing this calculation reveals that G_O2_ of the rectal gill increases from 0.09 in normoxia to 0.21 μl O_2_/min/kPa during 5.3 kPa. Thus, dragonfly nymphs are indeed increasing their physiological G as the water O_2_ concentration falls, indicating that they enhance diffusive O_2_ transfer through both hemolymph PO_2_ and O_2_ conductance in tandem with convective O_2_ transfer in order to maintain OEE and metabolic rates during hypoxia.

### Dragonfly nymph rectal gill conductance relative to other animals

Comparing the whole-gill O_2_ conductance of dragonfly nymphs with those measured from other gilled water-breathers reveals that the converted dragonfly value of 0.01 ml O_2_/min/kg/Torr lies well within the range of those seen in water-breathing fish (e.g. Baumgarten-Schumann and Piiper, 1968; Fernandes et al., 2012; Fisher et al., 1969; Holeton, 1970; Holeton, 1971; Hughes, 1972; Randall et al., 1967). This indicates that for a given unit of time and partial pressure gradient, O_2_ is as equally permeable across the dragonfly rectal gill as it is across the fish gill, and that the secondary evolution of water-breathing organs in these insects does not appear to have negatively affected their ability to extract O_2_. However, this may not be the case regarding the permeability of carbon dioxide (CO_2_). It is widely accepted that due to the difference in diffusivity and solubility of O_2_ and CO_2_ in water/tissue (physical parameters that are independent of the animal), the CO_2_ conductance is generally 20-fold greater than that of O_2_ (Dejours, 1989; Randall et al., 1967). Although the CO_2_ conductance of dragonfly nymph rectal gill has yet to be measured, it can be estimated based on measurements of metabolic rate, respiratory exchange ratio (RER), and hemolymph PCO_2_ by:

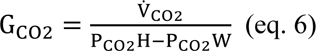

whe re V̇_CO2_ is CO_2_ production rate (calculated from V̇_O2_ and RER), PCO_2_H is hemolymph PCO_2_, and PCO_2_W is air-saturated water PCO_2_. The above calculation shows that assuming a dragonfly nymph has an RER of 0.73 (Harter et al., 2017) and PCO_2_H of 0.9 kPa (Lee et al., 2018), they would have a G_CO2_ of 1.1 μl CO_2_/min/kPa, a value that is just 13-fold higher than G_O2_ and noticeably lower than the 20-fold difference predicted from Krogh’s coefficient of O_2_ and CO_2_ in water/tissue. A lower G_CO2_ in dragonfly nymphs compared with other animals may present a mechanism for the elevated hemolymph PCO_2_ and total CO_2_ measured previously (Lee et al., 2018). A reduced CO_2_ permeability across the rectal gill would inhibit CO_2_ excretion in dragonfly nymphs, leading to an accumulation of total CO_2_ in the hemolymph, both as gaseous dissolved CO_2_ (PCO_2_) and hydrated bicarbonate (HCO_3_^-^). A more accurate measurement of G_O2_ and G_CO2_ will be necessary to test whether dragonflies do indeed have reduced CO_2_ permeability across their rectal gill.

### Conclusions

The observed ability of the tidally-breathing dragonfly nymphs to sustain their metabolic rates and O_2_ extraction efficiency during progressive hypoxia cannot be solely explained by an increase in ventilation frequency, and other parameters must also be changing to enhance O_2_ uptake from the water. Measuring the ventilatory and respiratory parameters of implanted nymphs exposed to progressive hypoxia reveals that the data from implanted nymphs are in excellent agreement with those of the control group. Empirical results and model simulations show that a significant reduction in hemolymph PO_2_ together with increased ventilation frequency can sustain the PO_2_ diffusion gradient and prevent OEE from falling during mild hypoxia, but not during severe hypoxia. An increase in O_2_ conductance is needed to sustain OEE at extremely low water PO_2_ in the model, and is indeed empirically observed in the dragonfly nymphs that increase their whole-gill O_2_ conductance during hypoxia. Lastly, a comparison of O_2_ and CO_2_ conductance of dragonfly nymphs to those of other water-breathers suggests that the rectal gill is less permeable to CO_2_, potentially explaining the elevated hemolymph CO_2_ content measured previously.

## Author contributions

D.J.L.: conceptualization, data curation, formal analysis, investigation, methodology, validation, visualization, writing-original draft, writing-review and editing; P.G.D.M.: conceptualization, funding acquisition, methodology, supervision, writing-original draft, writing-review and editing

## Acknowledgements

We thank the UBC Botanical Garden and Dolph Schluter for allowing us to collect dragonfly nymphs from their ponds, and the reviewers for their valuable comments and suggestions on the draft.

## References

Bäumer, C., Pirow, R. and Paul, R. J. (2000). Respiratory adaptations to running-water microhabitats in mayfly larvae *Epeorus sylvicola* and *Ecdyonurus torrentis*, Ephemeroptera. Physiological and Biochemical Zoology 73, 77–85.

Baumgarten-Schumann, D. and Piiper, J. (1968). Gas exchange in the gills of resting unanesthetized dogfish (*Scyliorhinus stellaris*). Respiration physiology 5, 317–325.

Birrell, J. H., Verberk, W. C. and Woods, H. A. (2023 preprint). Consistent Differences in Tissue Oxygen Levels Across 15 Insect Species Reflect a Balance between Toxicity and Asphyxiation. Available at SSRN 4592225.

Brainerd, E. L. and Ferry-Graham, L. A. (2005). Mechanics of respiratory pumps. Fish Physiol. 23, 1–28.

Brodersen, K. P., Pedersen, O., Lindegaard, C. and Hamburger, K. (2004). Chironomids (Diptera) and oxy-regulatory capacity: an experimental approach to paleolimnological interpretation. Limnology and Oceanography 49, 1549–1559.

Dejours, P. (1989). From comparative physiology of respiration to several problems of environmental adaptations and to evolution. J. Physiol. 410, 1–19.

Feder, M. E. and Burggren, W. W. (1985). Cutaneous gas exchange in vertebrates: design, patterns, control and implications. Biol. Rev. 60, 1–45.

Fernandes, M. N., da Cruz, A. L., da Costa, O. T. F. and Perry, S. F. (2012). Morphometric partitioning of the respiratory surface area and diffusion capacity of the gills and swim bladder in juvenile Amazonian air-breathing fish, *Arapaima gigas*. Micron 43, 961–970.

Fisher, T., Coburn, R. and Forster, R. (1969). Carbon monoxide diffusing capacity in the bullhead catfish. J. Appl. Physiol. 26, 161–169.

Galic, N., Hawkins, T. and Forbes, V. E. (2019). Adverse impacts of hypoxia on aquatic invertebrates: A meta-analysis. Science of the total environment 652, 736–743.

Harter, T. S., Brauner, C. J. and Matthews, P. G. (2017). A novel technique for the precise measurement of CO_2_ production rate in small aquatic organisms as validated on aeshnid dragonfly nymphs. J. Exp. Biol. 220, 964–968.

Hills, B. and Hughes, G. (1970). A dimensional analysis of oxygen transfer in the fish gill. Respiration physiology 9, 126–140.

Hitchman, M. L. (1978). Measurement of dissolved oxygen. New York: John Wiley & Sons.

Holeton, G. F. (1970). Oxygen uptake and circulation by a hemoglobinless Antarctic fish (*Chaenocephalus aceratus lonnberg*) compared with three red-blooded Antartic fish. Comp. Biochem. Physiol. 34, 457–471.

Holeton, G. F. (1971). Oxygen uptake and transport by the rainbow trout during exposure to carbon monoxide. J. Exp. Biol. 54, 239–254.

Hughes, G. and Morgan, M. (1973). The structure of fish gills in relation to their respiratory function. Biol. Rev. 48, 419–475.

Hughes, G. and Saunders, R. (1970). Responses of the Respiratory Pumps to Hypoxia in the Rainbow Trout (Salmo Gairdnert). J. Exp. Biol. 53, 529–545.

Hughes, G. M. (1972). Morphometrics of fish gills. Respiration physiology 14, 1–25.

Lee, D. J., Gutbrod, M., Ferreras, F. M. and Matthews, P. G. D. (2018). Changes in hemolymph total CO2 content during the water-to-air respiratory transition of amphibiotic dragonflies. Journal of Experimental Biology 221, jeb181438.

Lee, D. J. and Matthews, P. G. (2024). Oxygen extraction efficiency of the tidally-ventilated rectal gills of dragonfly nymphs. Proc. Roy. Soc. B 291, 20231699.

Maina, J. N. (2002). Structure, function and evolution of the gas exchangers: comparative perspectives. J. Anat. 201, 281–304.

Malison, R. L., DelVecchia, A. G., Woods, H. A., Hand, B. K., Luikart, G. and Stanford, J. A. (2020a). Tolerance of aquifer stoneflies to repeated hypoxia exposure and oxygen dynamics in an alluvial aquifer. J. Exp. Biol. 223, jeb225623.

Malison, R. L., Ellis, B. K., DelVecchia, A. G., Jacobson, H., Hand, B. K., Luikart, G., Woods, H. A., Gamboa, M., Watanabe, K. and Stanford, J. A. (2020b). Remarkable anoxia tolerance by stoneflies from a floodplain aquifer. Ecology 101, e03127.

Perry, S., Jonz, M. and Gilmour, K. (2009). Oxygen sensing and the hypoxic ventilatory response. In Fish physiology, vol. 27 eds. J. G. Richards A. P. Farrell and C. J. Brauner), pp. 193–253: Academic Press.

Piiper, J. and Scheid, P. (1972). Maximum gas transfer efficacy of models for fish gills, avian lungs and mammalian lungs. Respiration physiology 14, 115–124.

Pinder, A. W. and Feder, M. E. (1990). Effect of boundary layers on cutaneous gas exchange. J. Exp. Biol. 154, 67–80.

Porteus, C., Hedrick, M. S., Hicks, J. W., Wang, T. and Milsom, W. K. (2011). Time domains of the hypoxic ventilatory response in ectothermic vertebrates. J. Comp. Physiol. B 181, 311–333.

R Core Team. (2022). R: A language and environment for statistical computing. R Foundation for Statistical Computing, Vienna, Austria.

Rahn, H. (1966). Aquatic gas exchange: theory. Respir. Physiol. 1, 1–12.

Randall, D., Holeton, G. and Stevens, E. D. (1967). The exchange of oxygen and carbon dioxide across the gills of rainbow trout. J. Exp. Biol. 46, 339–348.

Rantin, F., Kalinin, A., Glass, M. and Fernandes, M. (1992). Respiratory responses to hypoxia in relation to mode of life of two erythrinid species (*Hoplias malabaricus* and *Hoplias lacerdae*). J. Fish Biol. 41, 805–812.

Robertson, H. T. (2015). Dead space: the physiology of wasted ventilation. Eur. Respir. J 45, 1704–1716.

Scott, G. R., Wood, C. M., Sloman, K. A., Iftikar, F. I., De Boeck, G., Almeida-Val V. M. and Val, A. L. (2008). Respiratory responses to progressive hypoxia in the Amazonian oscar, Astronotus ocellatus. Resp. Physiol. Neurobi. 162, 109–116.

Thomas, J. and Gilmour, K. (2012). Low social status impairs hypoxia tolerance in rainbow trout (*Oncorhynchus mykiss*). J. Comp. Physiol. [B] 182, 651–662.

Ultsch, G. R. (1996). Gas exchange, hypercarbia and acid-base balance, paleoecology, and the evolutionary transition from water-breathing to air-breathing among vertebrates. Palaeogeogr. Palaeoclimatol. Palaeoecol. 123, 1–27.

Weibel, E. R. (1984). The pathway for oxygen: structure and function in the mammalian respiratory system. Cambridge, US: Harvard University Press.

